# A simple methods for obtaining a multivalent protein with a variable number of binding sites and estimating its binding parameters

**DOI:** 10.1101/2023.11.28.569006

**Authors:** Alisa Mikhaylina, Natalia Lekontseva, Albina Khairetdinova, Nelly Ilyina, Vitalii Balobanov

## Abstract

The study of natural and the design of artificial multivalent proteins is a promising field of molecular biology. Working with such proteins is much more difficult than with their monovalent analogues. In this paper, we show how using a ring of heptameric Sm-like protein as a scaffold, it is possible to create a multivalent protein with a different number of binding sites. This is an urgent task for the study of multivalent and multicenter protein-protein interactions. The method of analysis used in the work allows us to evaluate the stoichiometry and the dissociation constant of complexes of artificial chaperone with a non-native protein. It is shown that for reliable binding of non-native αLA, its interaction with several apical domains of GroEL is necessary. At the same time, the dissociation constant of such a complex does not significantly change with an increase in the number of binding domains in the oligomer. Up to 4 αLA molecules can be attached to the complete heptameric ring of apical domains. The proposed methods have a good cost-to-result ratio and can be applied to the study and design of other new proteins.

## 1. Introduction

Multidomain proteins that have multiple binding sites are widespread in nature. First of all, these are various antibodies of the IgE and IgM classes. IgE has four antigen binding sites and IgM has ten. No less remarkable are the multivalent hormone/receptor complexes [1,2]. Chaperone molecules also have many binding sites [3]. The multivalency of the protein can increase binding affinity, avidity and specificity compared to monovalent analogues [4]. A multivalent interaction can be strong even if its individual bonds are weak [5,6]. No less interesting is that multivalency significantly increases the sensitivity of the protein-ligande interaction to external conditions, which leads to a hypersensitive and highly nonlinear dependence of the binding strength on parameters such as temperature, pH or components concentration [7,8]. In biological systems, low-affinity proteins are generally grouped together to achieve the required binding strength. From an evolutionary point of view, it seems convenient to combine several single interactions with low affinity to produce a collectively stronger result. In addition, tunable multivalent receptor–ligand interactions provide differential responses to biological signals.

As you can see from the above, confusion in understanding can be caused by the fact that the binding of several identical ligands by one protein and the binding of the same ligand by several binding sites in the literature are called the same - multivalent binding. Therefore, before continuing my story, it is worth focusing on the differences between these variants of multivalency. So: a typical IgG antibody binds two ligands at two independent binding sites and is therefore bivalent. The binding of a non-native protein to a chaperone that involves one ligand and several chaperone subunits can also be called multivalent [9]. These concepts of multivalency should be clearly distinguished. In an experiment, multivalent binding will be manifested by anomalous behavior of association/dissociation kinetic curves. In the case of independent binding of each ligand to each binding site, there will be no fundamental difference from the simple model (example immunoglobulin) [10]. There will be a difference only in the effective concentration and stoichiometry of the complex formed. Significant differences will occur when one ligand binds to multiple binding sites (example chaperone). The nonlinearity of the binding force and the hyperselectivity mentioned above will also manifest here [11].

### Determination of protein-protein interaction characteristics

The main interest in studying the interaction of multivalent proteins is to determine the parameters of their interaction with ligands. Kinetic and equilibrium characteristics can be determined by various methods. We will now describe the main experimental methods used for this.

Currently, microscale thermophoresis (MST), isothermal titration calorimetry (ITC), biolayer interferometry (BLI) and surface plasmon resonance (SPR) are mainly used to determine biomolecular interactions. Each method has its advantages and disadvantages. The microscopic thermophoresis method (MST) is based on measuring of changes in the fluorescence of molecules in microscopic temperature gradients, which are created using an infrared wavelength laser (IR laser) [12]. Any change in thermophoretic properties upon addition of a partner molecule is manifested as a change in fluorescence intensity. The normalized fluorescence is determined, which reflects the change in the concentration of molecules. Measuring these changes allow to quantify the affinity of interacting partners. Advantages of MST is small sample size, sample is not immobilized, low sample consumption, ability to measure complex mixtures, wide size range of interactants. Disadvantages of this technique is requiring of ligand fluorescent labelling, absence of kinetic and concentration analysis, high cost of ownership, high cost of experiment.

Isothermal Titration Calorimetry (ITC) is a technique used to quantify the thermodynamics of biomolecular interactions [13]. ITC measures the heat evolved during molecular interactions when a ligand is titrated with an analyte. The information obtained is the power required to maintain a constant temperature between the sample cell and the reference cell. This normalizing ability is measured as a function of time and used to determine the heat generated when a ligand interacts with its partner. Advantages of ITC is ability to determine multiple thermodynamic binding parameters (association constant, binding enthalpy and stoichiometry,) in a single experiment, absence of binding partners modification necessity, automated experimental process. Disadvantages of this technique is large sample quantity and volume, non-covalent complexes may exhibit small binding enthalpies as signal is proportional to the binding enthalpy, a long time for experiment.

Surface plasmon resonance (SPR) is a label-free technique that allows to quantify binding between biomolecules [14]. The SPR signal is based on changes in the refractive index at the surface of a gold sensor chip as an analyte flows in a microfluidic channel and binds to a ligand immobilized on the sensor chip. Monitoring the change in the SPR signal over time produces a sensorgram, a plot of the binding response versus time. Fitting the sensorgram data to a suitable kinetic binding model allows for the calculation of kinetic parameters such as the association (ka) and dissociation (kd) rate constants. Advantages of SPR is label-free detection, high throughput capabilities, high sensitivity with reproducible results, relatively small sample consumption. Disadvantages of this technique is high cost of ownership, ongoing fluidic maintenance, high cost of experiment, requiring immobilization of one of the binding partners.

Bio-Layer Interferometry (BLI) is an optical analytical technique used to quantify biomolecular interactions [15]. It analyzes the interference pattern of white light reflected the immobilized protein layer at the tip of the fiber-optic sensor and an internal reference layer. The binding of solution molecules to the biosensor tip causes a shift in the interference pattern. This shift is measured to identify, quantify, and characterize proteins and other biomolecules in solution. Advantages of BLI is label-free detection, reference channel-free detection, fluidic-free system means less maintenance needed. Disadvantages of this technique is immobilization of ligand to surface of tip required and low sensitivity (100-fold lower sensitivity of detection compared to SPR).

The theoretical description of the kinetics and thermodynamics of processes occurring during polyvalent interaction is complex. The theory is discussed in sufficient detail in the works [10,16]. Theoretical models can be used to obtain thermodynamic interaction parameters based on experimental data obtained using the methods described above. However, there is a rather serious problem - the more complex model leads more ambiguous in experimental data describing. The researcher needs to clearly understand the required limit of detail in describing the process. A simple comparison of several protein variants with each other is often required [17]. Determining the equilibrium dissociation constants of complexes also rarely requires great accuracy. At the same time, the excessive complexity of the used mathematical model can worsen the obtained result. In our work, we tried to simplify as much as possible the experimental methods used and the processing of results. The reader will be able to assess how far we succeeded in this.

### Artificial chaperone ADGroEL_SacSm

The object of our study is the artificial chaperone ADGroEL_SacSm. Like the GroEL chaperone, the Apical Domain (ADGroEL) of which is part of ADGroEL_SacSm [18]. The binding of non-native proteins by chaperones is described as multivalent [19].That is, when the bound protein interacts with many sites on the chaperone. These interactions are described as nonspecific and individually relatively weak. To evaluate the details of non-native proteins binding by our chaperone, we undertook this study. We developed and applied two simple methods. The first method is to obtain chaperones with different numbers of binding sites. The second is to assess stoichiometry and binding strength.

The principle of obtaining ADGroEL_SacSm that we used was the ADGroEL attachment to the oligomeric scaffold of the SacSm protein.

### SacSm

The basis for our design is annular homoheptameric Sm-like protein of the archaea *Sulfolobus acidocaldarius*. These proteins are highly resistant to denaturants and are hyperthermostable. Analysis of the structure of SacSm shows that the simplest way to attach the target domain to it is to connect it by a flexible linker [18] to the N or C terminus. Similar Sm-like proteins are present in bacteria, archaea and eukaryotic organisms [20,21]. Often these are RNA-binding proteins. They are also often attributed to the role of RNA chaperones. There are proteins with the number of monomers in the ring from 5 to 8. Many of them have high structural stability [22,23]. They are an excellent basis for work similar to ours one.

### GroEl and ADGroEL

GroEL is a well-studied protein chaperone. Structurally, it is a dodecamer of 2 rings of 7 subunits each. Each subunit consists of three domains: Apical, Intermediate and Equatorial. The ADGroEL plays a major binding role. In addition, back in 1996, it was shown that the monomeric ADGroEL, separated from the rest of the structure, retains chaperone function [24]. Structurally, the ADGroEL is anchored by both ends of its polypeptide chain. Both ends are located in the Equatorial domain. At the same time, the interaction of the ADGroEL with each other is quite weak and they change mutual orientation when ATP is bound by the Equatorial domain.

### ADGroEL_SacSm

We combined SacSm and ADGroEL to create a new chaperone ADGroEL_SacSm. The created protein showed good ability to bind non-native proteins and prevent its aggregation when heated. At the same time, a monomeric ADGroEL in our experiments did not exhibit such properties [18].

The high stability of SacSm allows it to maintain its structure even in high concentrations of urea [23]. We also previously demonstrated effective renaturation of the ADGroEL attached to a scaffold of the SacSm protein. When renatured from the fully unfolded state (high concentration of guanidine hydrochloride), the significant difference in stability between the proteins parts results in two-step folding. The first step is the folding of the SacSm and the heptameric ring formation. The second step is the folding of added ADGroEL.

### The problem definition

As mentioned above, the heptomer ADGroEL_SacSm is able to bind non-native proteins, but the monomeric ADGroEL is not. The question arises: how many ADGroELs collected together are enough for a strong binding? For full-size GroEL, the answer to this question was given quite some time ago in article [3]. As has been shown, the minimum element required for binding is two ADGroELs located next to each other. But what about our protein? In our protein, the mutual mobility of domains is higher than in GroEL, and in addition, in the work mentioned above, the interaction was switched off by mutations, which does not exclude additional interactions. These questions led us to the idea of using a system for varying the number of domains in our oligomer.

The significant difference in stability of the base and the attached domain allows the following manipulations. If when the hybrid protein is completely unfolded by a high concentration of guanidine hydrochloride, the SacSM (without attached ADGroELs) is added, then upon further folding it will be included in the scaffold ring. This inclusion will be random and proportional to the ratio of the ADGroEL_SacSm and empty SacSM. In such a situation, the output will be proteins with different numbers of ADGroELs. The remarkable thing about this system is its versatility and applicability with other attached domains, provided they are effectively renatured.

To assess the stoichiometry of the resulting complexes and the strength of binding between their constituent proteins, we developed a technique based on metal-affinity chromatography and quantitative analysis of its results by SDS gel electrophoresis.

## 2. Materials and Methods

### Purification of the Fusion Protein ADGroEL_SacSm

The pET-22b vector carrying the ADGroEL_SacSm fusion protein gene was used for transformation in *E. coli* BL21(DE3)/pRARE to express the protein. *E. coli* strain BL21(DE3) cells were preliminarily co-transformed with the pRARE, which carried rare-codon tRNA genes (AUA, AGG, AGA, CUA, CCC, and GGA) to enhance the expression of fusion proteins, including an archaeal SacSm part that contained codons rarely used in *E. coli*.

The transformants were grown in an LB medium in the presence of ampicilline (100 μg/mL) and chloramphenicol (10 μg/mL), at 37 °C with an agitation of 180 rpm. Protein expression was induced at OD600nm = 0.6–0.8 o.u. with the addition of IPTG at a final concentration of 0.5 mM. The bacteria were harvested with centrifugation 3 h after in-duction.

Cell pellets were suspended in a solution containing 500 mM NaCl, 50 mM Tris-HCl (pH 8.0), 10 mM imidazole, 5 mM β-mercaptoethanol, and 6M Guanidine-HCl. Cells were disrupted by sonication at 4°C. Cell debris was removed by centrifugation at 15,000× g for 30 min at 4°C. Supernatant was loaded onto a Ni-NTA agarose (GE Healthcare, Uppsala, Sweden) column equilibrated with a solution containing 500 mM NaCl, 50 mM Tris-HCl (pH 8.0), and 10 mM imidazole and 8M Urea. ADGroEL_SacSmAP were eluted using a step gradient of imidazole (40 mM and 150 mM) in a solution containing 500 mM NaCl and 50 mM Tris-HCl (pH 8.0) and 6M Urea. Fractions containing the protein were collected and purified using Q-Sepharose equilibrated with a solution containing 500 mM NaCl, 50 mM Tris-HCl (pH 8.0). Fractions containing the protein were concentrated and dialyzed against a solution containing 150 mM NaCl and 50 mM Tris-HCl (pH 8.0). The final purification step was size-exclusion chromatography on the Superdex 75 resin equilibrated with a solution containing 150 mM NaCl and 50 mM Tris-HCl (pH 8.0). Fractions containing the protein were concentrated and dialyzed against a solution containing 150 mM NaCl and 50 mM Tris-HCl (pH 8.0).

### Expression and purification of SacSm

Competent BL21(DE3)/pLacIRARE *E. coli* were transformed with pProExHTb SacSm and grown in LB media (100 mg/ml ampicillin and 10 mg/ml chloramphenicol) at 37°C to an optical density at 600 nm (OD600) of ≈0.8. Then, overexpression of SacSm was induced by adding IPTG to a final concentration of 0.5 mM. After overnight incubation at 20°C, cells were centrifuged at 14000× g for 30 min.

The cell pellet from the previous step was resuspended in Lysis Buffer (20 mM sodium phosphate buffer, pH 8.0, 0.5 M NaCl, 10 mM imidazole, 1 mM PMSF, 1 mM DTT, 0.1% Triton X-100) and disrupted by sonication (Fisher Scientific, USA). The extract was cleared by centrifugation at 14,000g for 30 min, then the supernatant was heated at 70°C for 20 min and denatured *E. coli* proteins were removed by centrifugation (14,000g, 40 min, 4°C).

SacSm-containing protein sample was loaded on a Ni-NTA Agarose (Qiagen) column equilibrated with 20 mM sodium phosphate buffer, pH 8.0, 0.2 M NaCl, 10 mM imidazole.

The His-tagged SacSm protein was eluted by applying a linear gradient of imidazole, from 10 mM to 250 mM. To proteolytically remove the His-tag SacSm-containing fractions were combined, concentrated and incubated with TEV protease at a 1:100 mass ratio of TEV:SacSmAP. To remove TEV protease from the sample, the solution was heated for 20 min at 65°C, then cleared by centrifugation (14,000g, 40 min, 4°C) and re-applied to the Ni-NTA Agarose to remove cleaved His-tags. The flow-through of SacSmAP was concentrated and applied to Superdex75, equilibrated with 50 mM Tris-HCl, pH 8.0, 300 mM NaCl, 1 mM DTT.

### Obtaining hetero-oligomers by the denaturation/renaturation

The schematic diagram of heterooligomers obtaining is shown in Figure 1. ADGroEL_SacSm and SacSm proteins were mixed in different molar ratios. Dry Guanidine hydrochloride was added to the protein solution up to saturation. Then a solution of 20 mM Hepes-NaOH pH 7.5, NaCl 200 mM was gradually added to the protein solution with 30 minutes incubation at each step until Guanidine hydrochloride concentration achieved 1M. The resulting mixture was applied to a chromatography column with Ni-Focurose metal chelate resin (Elabscience, China). The column was washed with buffer contained 20 mM Hepes-NaOH pH 7.5, NaCl 200 mM. The target protein was eluted from the Ni-Focurose column with the same solution with the addition of 500 mM Imidazole.

**Figure 1.**
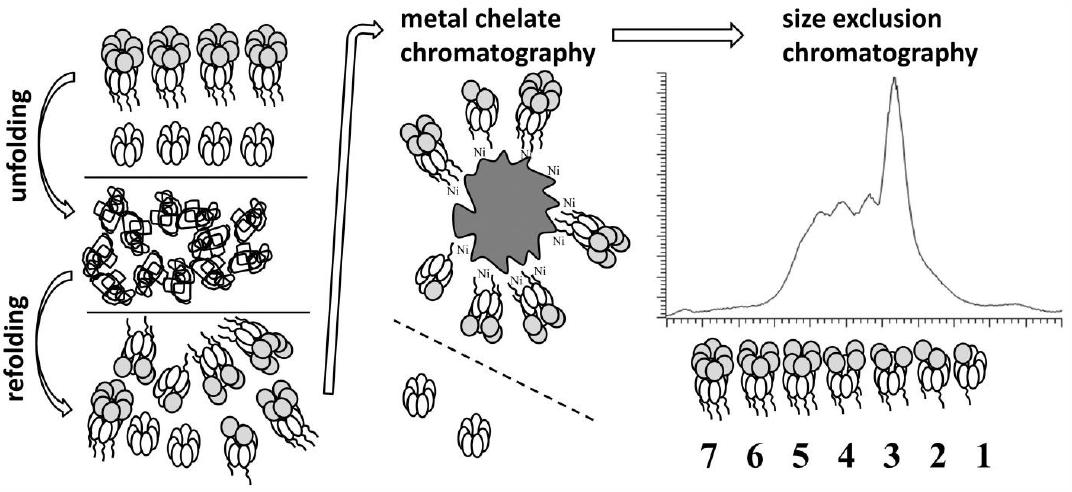
A scheme for obtaining heterooligomers with a different number of binding domains. A protein with a binding domain has his-tag and protein without such domain does not have it. After collaborative refolding, proteins having a binding domain are purified on a metallochelate resin and separated by size exclusion chromatography.

The separation of hybrid forms of proteins was carried out with size-exclusion chromatography using Superdex 200 Increase 10/300 GL column. Chromatography was performed on the AKTA Basic FPLC system (Amersham, Sweden). The volume of protein injected was 300 μl, with a total protein concentration of 3-4 mg/ml. Working solution contained 20 mM Hepes-NaOH pH 7.5, NaCl 200 mM. The flow rate was 0.4 ml/min. Proteins containing varying amounts of ADGroEL were separated according to their molecular weight. The volume of each fraction was 200 μl.

### Non-native protein binding study

The schematic diagram of non-native protein binding experiment is shown in Figure 2. We used denatured alpha-Lactalbumin (αLA) (Sigma, USA) as a model substrate for protein binding assay. αLA was in solution contained 20 mM Hepes-NaOH pH 7.5, NaCl 200 mM, Urea 6M, beta-mercaptoethanol 10mM before use. αLA was added to a solution of ADGroEL_SacSm/SacSm hetero-oligomers in a ratio of 7 αLA molecules per one ring of ADGroEL_SacSm/SacSm in a buffer solution contained 20 mM Hepes-NaOH pH 7.5, NaCl 200 mM, beta-mercaptoethanol. In the presence of a reducing agent, αLA is unable to assume its native structure. The protein solution was incubated for 10 minutes followed by applying to Ni-Focurose and unbound αLA was washed off with the same buffer solution. Proteins bound to the Ni-Focurose were eluate with a buffer solution 20 mM Hepes-NaOH pH 7.5, NaCl 200 mM, Imidazole 500 mM and analyzed by SDS-PAGE.

**Figure 2.**
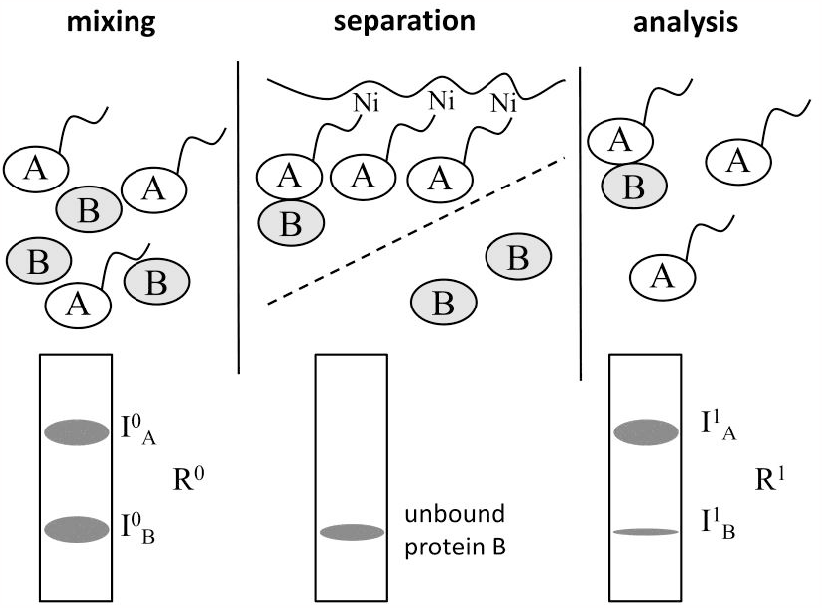
The study design for investigating protein-protein binding. Protein A has his-tag and protein B does not. After co-incubation, protein B not bounded to A is separated by metallochelate chromatography. Each stage was analyzed by SDS-PAGE.

### SDS-PAGE

SDS-PAGE was used to analyze the purity and quantitative characterization of the hybride ADGroEL_SacSm/SacSm proteins.

SDS polyacrylamide gel electrophoresis (SDS-PAGE) was performed according to the Laemmli method with some modifications.

The separating gel contained: 15% acrylamide (AA:MBA ratio is 39:1), 0.375 M Tris-HCl, pH 8.8, 0.1% SDS. 0.1% TEMED and 0.1% APS were added for polymerization. The stacking gel contained: 6% acrylamide (AA:MBA ratio is 39:1), 0.125 mM

Tris-HCl, pH 6.8, 0.1% SDS. 0.1% TEMED and 0.1% APS were added for polymerization.

The gel was carried into the gel plates measuring 7 × 8 cm and 0.75 mm thick.

Samples were prepared by adding Laemmli sample buffer in an amount of 1/5 of the sample volume and boiled for 5 min. Laemmli sample buffer contained 0.3 M Tris-HCl, pH 6.8, 10% SDS, 25% β-mercaptoethanol, 30% glycerol, 0.1% BromPhenol Blue.

The running buffer used was Tris/Glycine/SDS and contained 25mM Tris, 192 mM Glycine, 0.1% SDS. Gels were run in the “Mini Protean II” electrophoresis cell from Bio-Rad (USA) at 200 V until the dye front reached the bottom of the gel.

After electrophoresis, gels were stained in 0.05% Coomassie G-250 solution in 30% Ethanol and 10% Acetic Acid. The background color was washed off by boiling in 5% Acetic Acid.

### SDS-PAGE images analysis and calculation of equilibrium dissociation constants

To quantitatively analyze the ratios of ADGroEL_SacSm and SacSm in hetero-oligomers and analyze the binding of αLA, the color intensities of the corresponding bands on the SDS-PAGE images were determined. The determination was carried out using the TotalLab Software (nonlinear dynamics, UK). The color intensity of the corresponding bands was calculated after subtracting the background color intensity. After this, considering the color intensity to be proportional to the protein mass in the band and taking into account its molecular weight, we can introduce the following relations:

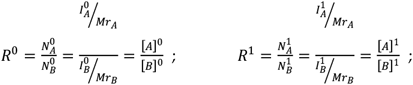

Since protein A fully binds to the resin and protein B binds only as part of the complex, then:

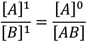

For the simplest variant of determining the dissociation constant, we will express the necessary concentration values through the relations that we have introduced

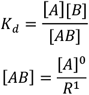

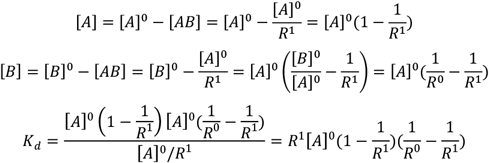

Thus, knowing the total concentration of protein A and determining the ratio of components before and after the separation of unbound protein B, we can estimate the dissociation constant of the resulting complex.

## 3. Results and Discussions

### Preparation of hetero-oligomers

Hetero-oligomers were obtained according to the method described in the Materials and Methods section. Different ratios of ADGroEL_SacSm and SacSm were tested. A change in this ratio expectedly leads to a shift in the distribution of proteins towards more or less ADGroEL_SacSm in the hetero-oligomer. With a starting ADGroEL_SacSm / SacSm ratio of 1/3, the distribution allowed us to obtain almost all variants in an acceptable quantity. We separated the mixture of the resulting hetero-oligomers into fractions by size-exclusion chromatography (fig.3A). Separation is more efficient for hetero-oligomers with small amount of ADGroEL domains, despite the same molecular weight step (per molecular weight of one ADGroEL), this is due to a smaller percentage change in the hydrodynamic size with a larger number of domains. Electropherogram patterns of the obtained fractions analysis made it possible to calculate the ADGroEL_SacSm / SacSm ratios in the obtained hetero-oligomers (fig.3B and table1). In general, the method of obtaining and analyzing is quite simple and intuitive. Its disadvantages include the inability to control the relative position of the ADGroEL on the ring. When transferring the technique to other proteins, their renaturation ability will also be a limitation.

**Table 1.** Calculation of the ADGroEL_SacSm / SacSm ratios in the obtained hetero-oligomers.

| Fraction number<br>(see fig.3) | Color intensity of<br>ADGroEL_SacSm<br>bands | Color<br>intensity<br>of<br>SacSm<br>bands | ADGroEL_SacSm /<br>SacSm molar ratio | ADGroEL_SacSm<br>number per ring |
| --- | --- | --- | --- | --- |
| 1 | 418449 | - | - | - |
| 2 | 600442 | 602505 | 0.28 | 1.6 |
| 3 | 1818168 | 879939 | 0.59 | 2.6 |
| 4 | 2942202 | 806784 | 1.04 | 3.6 |
| 5 | 3202075 | 197672 | 4.61 | 5.8 |

**Figure 3.**
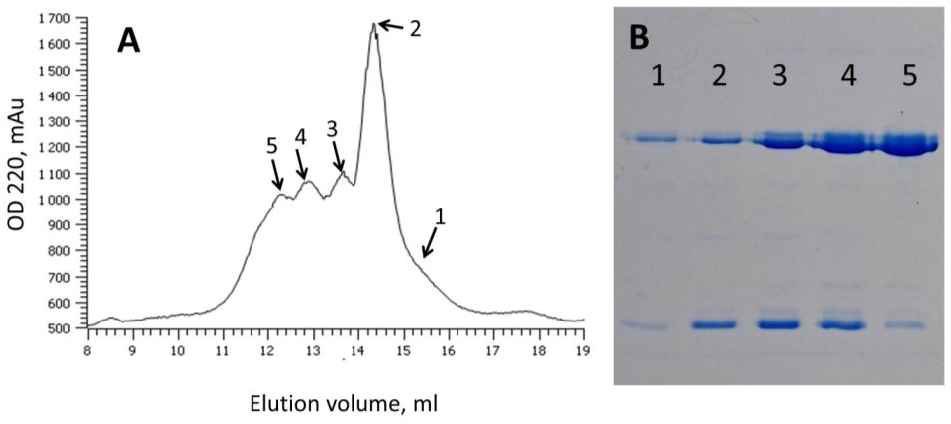
Results of separation of ADGroEL_SacSm / SacSm heterooligomers with different numbers of binding domains by size-exclusion chromatography. Chromatogram - A and SDS-PAGE analysis of the corresponding fractions - B. The quantitative assessment is given in table 1.

### Non-native protein binding assay

We used αLA as a model of non-native protein. In the presence of a reducing agent, αLA is unable to assume its native structure[].We obtained complexes of chaperones and non-native proteins by their joint incubation. To evaluate the binding of non-native proteins, we separate bound and unbound non-native proteins on a Ni-Focurose metal chelate resin. αLA does not bind to this resin. ADGroEL_SacSm and obtained hetero-oligomers have a His-tag and therefore can binding to resin.

After washing the not bound proteins, only αLA bound to ADGroEL_SacSm remains bound to Ni-Focurose. The results show that hetero-oligomers containing only 2 ADGroEL_SacSm subunits do not bind to non-native αLA (fig.4). This experiment is also a control for the binding of αLA to the Ni-Focurose and to the SacSm oligomer, which serves as the scaffold of the hetero-oligomer. In our case, such binding does not occur. αLA reliably binds to hetero-oligomers with three or more ADGroEL_SacSm in their composition (fig.4 and table2).

**Figure 4.**
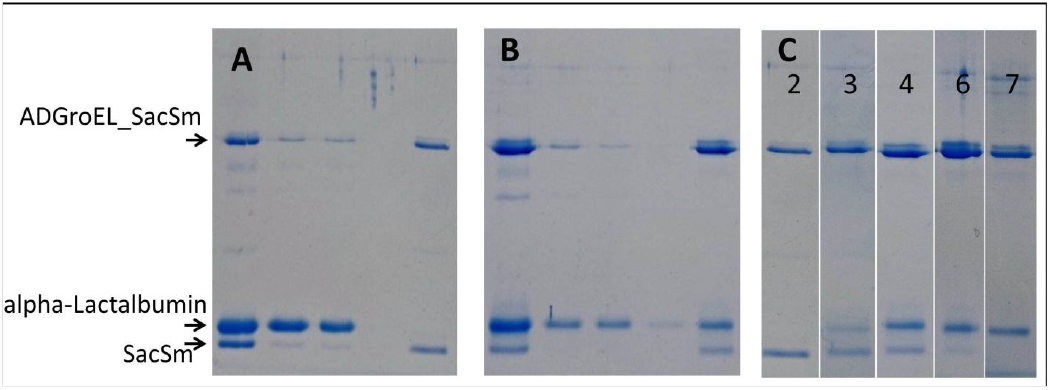
Analysis of the binding of non-native proteins by chaperone. The first line in fields A and B is the initial mixture, the last line is after the separation of the unbound protein. Intermediate lines - resin washing. Field C is a combination of lines after separation of an unbound protein. The number of binding domains in heterooligomer indicated on the corresponding line.

**Table 2.** Calculation of stoichiometry and dissociation constants based on the data in figure 4.

| ADGro-EL_SacSm num-<br>ber per ring | ADGro-EL_SacSm /αLA<br>molar ratio, ini-<br>tial | ADGro-EL_SacSm /αLA<br>molar ratio, in<br>complex | αLA mole-<br>cules per het-<br>ero-oligomer<br>ring | Kd<br>(M) |
| --- | --- | --- | --- | --- |
| 2 | 0.32 |  |  |  |
| 3 | 0.50 | 2.49 | 0.96 | $4 \cdot 10^{-6}$ |
| 4 | 0.72 | 2.14 | 1.75 | $3 \cdot 10^{-6}$ |
| 6 | 0.90 | 2.34 | 2.44 | $4 \cdot 10^{-6}$ |
| 7 | 0.57 | 1.57 | 4.41 | $3 \cdot 10^{-6}$ |

To solve the problem of assessing the stoichiometry of the resulting complex and the binding strength of the substrate protein, we used the same experimental approach. Of course, more precise methods could have been used, such as SPR or MST. But a rough estimate was enough for us. In addition, we wanted to obtain a simpler and cheaper experimental technique. And we got it.

The results obtained show that binding is detected only for a hybrid containing at least 3 ADGroEL in the ring. In this case, the binding stoichiometry is close to the ratio of one αLA per two ADGroEL. This can be explained by the binding of two adjacent ADGroELs of one αLA. Which correlates well with the data for a full-size GroEL from the article [3]. However, discrepancies with the literature data were also found. Thus, the stoichiometry of the interaction of full-length GroEL with non-native proteins is estimated as one non-native protein per one GroEL ring, that is, 7 ADGroELs. Our artificial chaperon binds 4 αLA molecules per complete ring. Most likely, this can be explained by the greater mobility of the ADGroELs relative to each other, which reduces the competition between the bound molecules.

We estimated the dissociation constant using the described above method. We calculated molar ratios of proteins before and after separation on the Ni-Focurose using color intensities of electropherogram bands. This approach is based on the assumption that the color intensity is proportional to the mass of the protein in this band. I agree, the assumption is quite rough. However, it is often necessary to estimate the dissociation constant from the order value and to compare the efficiency of interaction between different variants of molecules. Such accuracy is quite achievable for the applied method. And it is quite sufficient for comparing dissociation constants determined uniformly among themselves by the same technique.

The calculation results are shown in the table. In the calculation, we consider the concentration of ADGroEL_SacSm as the concentration of binding sites. The result shows that the dissociation constant practically does not change when the proportion of ADGroEL_SacSm in the hetero-oligomer changes.

The totality of the data obtained suggests the following picture: the interaction of a single ADGroEL with non-native proteins is quite weak. This prevents it from being detected by our methods. But such interaction is sufficient for it to exhibit chaperone activity, as follows from the literature data [24]. When several ADGroELs are located closely on a single base, non-native proteins can interact with several domains simultaneously and, accordingly, the stability of such a complex increase exponentially. Our model protein αLA is small and cannot interact with more than two ADGroELs, which is manifested by the absence of a change in the dissociation constant with an increase in the proportion of ADGroEL_SacSm in the hetero-oligomer. At the same time, its small size allows up to 4 molecules to be placed on the complete heptameric ADGroEL_SacSm ring.

## 5. Conclusions

Thus, the two methods presented in this article allowed us to identify important details of the functioning of the artificial chaperone ADGroEL_SacSm. Comparison of these methods with alternative approaches shows their advantages.

Varying the number of binding sites on a protein by assembling hetero-oligomers can significantly reduce the cost and time spent compared to alternative approaches. You can agree, obtaining an individual genetic construct for each variant and isolating each variant of the corresponding proteins is much more difficult.

The proposed method for estimating the stoichiometry of the resulting complexes and the strength of interaction in them is rather coarse. But if you compare our approach with alternative methods, it becomes clear that its accuracy is sufficient for many tasks. At the same time, it does not require expensive equipment and qualified staff. If greater accuracy is required, then our proposed approach may well be used as the first stage of searching for conditions for further experiments by other methods.

